# Human fallopian tube single-cell analysis reveals monocytic transcriptional and interaction changes in high-grade serous ovarian cancers

**DOI:** 10.1101/2023.07.14.549073

**Authors:** Joshua Brand, Marcela Haro, Xianzhi Lin, Stephanie M. McGregor, Kate Lawrenson, Huy Q. Dinh

**Author notes:** Correspondence: Huy Dinh.

## Abstract

Tumorigenesis for most high-grade serous ovarian cancers (HGSCs) likely initiates from fallopian tube (FT) epithelia. While epithelial subtypes have been characterized using single-cell RNA-sequencing (scRNA-Seq), heterogeneity of other cellular compartments and their involvement in tumor progression are poorly defined. Integrated analysis of human FT scRNA-Seq data and other relevant tissues, including HGSC tumors, revealed greater transcriptional diversity of immune and stromal cells. We identify an unprecedented abundance of monocytes in human FT myeloid cells across two independent donor cohorts. The ratio of macrophages to monocytes are relatively similar between benign FTs, ovaries, and adjacent normal tissues, but is significantly greater in tumor. FT-defined monocyte and macrophage signatures, cell-cell communication, and gene set enrichment analysis identified monocyte- and macrophage-specific ligand-receptor interactions and functional pathways in tumors and adjacent normal tissue. Further reanalysis of tumor scRNA-Seq from HGSC patients suggested different monocyte and macrophage subsets associated with neoadjuvant chemotherapy treatment. Taken together, our work provides evidence that an altered FT immune composition could inform early detection markers in HGSCs.

## Introduction

A lack of early detection markers for high-grade serous ovarian cancers (HGSCs) has contributed to patients’ poor clinical outcomes and stagnant survival rates over several decades (Lisio et al. 2019). Emerging molecular data show that the majority of these cancers likely arise from secretory cells at the fimbriated end of the fallopian tube (FT) (Labidi-Galy et al. 2017; Ducie et al. 2017; Eckert et al. 2016; Y. Lee et al. 2007; Lawrenson et al. 2019), emphasizing the importance of characterizing the FT tissue microenvironment to gain insight into the earliest stages of HGSC development. While significant efforts have been made to characterize the tumor microenvironment (TME) of HGSCs (Qian et al. 2020; Binnewies et al. 2021; Olalekan et al. 2021; Olbrecht et al. 2021; Xu et al. 2022; Hornburg et al. 2021) and benign FT epithelia (Dinh, Lin, et al. 2021; Hu et al. 2020; Ulrich et al. 2022) using single-cell transcriptomics (scRNA-Seq), there remains a critical need to connect the single-cell landscape of stromal and immune subsets of human FT to HGSC.

Limited data have been reported on the extended heterogeneity of human FT immune cell subsets and their cell-cell interactions. Previous studies used protein-panel-based technologies such as immunohistochemistry (IHC) (George, Milea, and Shaw 2012) and flow cytometry (Shaw et al. 2011; Ardighieri et al. 2014; S. K. Lee et al. 2015) to profile human FT immune subsets with limited markers for phenotyping beyond those for cell type identification (e.g. macrophages, DCs, and T/NK cells). The heterogeneity of T/NK, myeloid, and stromal cells and their cell-cell interactions in FT remain incomplete in published scRNA-Seq analyses (Dinh, Lin, et al. 2021; Ulrich et al. 2022). We therefore analyzed immune and stromal cell heterogeneity from scRNA-Seq of benign human FTs (Dinh, Lin, et al. 2021; Ulrich et al. 2022) and HGSC tumors and adjacent normal samples from >80 in-house and publicly available sources (Ulrich et al. 2022; Binnewies et al. 2021; Olalekan et al. 2021; Xu et al. 2022; Yu et al. 2022; Zhang et al. 2022; Qian et al. 2020), identifying 7 fibroblast/stromal, 6 T/NK and 7 myeloid cell subsets with distinct gene, pathway, and regulatory network signatures. Further examination of single-cell derived transcriptomic signatures revealed associations with molecular subtypes in bulk transcriptome data from The Cancer Genome Atlas (TCGA). Lastly, we show a significant macrophage-to-monocyte ratio shift in tumor compared to benign FT and transcriptional changes in FTs from women with *BRCA*1/2 mutations and HGSC patients treated with chemotherapy. These results provide a better understanding of human FT heterogeneity and its implications for HGSC outcomes.

## Results

### ScRNA-Seq analysis of non-epithelial cell compartment in human fallopian tubes

To characterize the cellular and molecular characteristics of the human FT tissue microenvironment, we reanalyzed the non-epithelial cell compartment from our previously published scRNA-Seq (Dinh, Lin, et al. 2021), which included 45,654 non-epithelial cells. Here, we focus on immune and stromal heterogeneity and compare our subsets to a recently published, independent scRNA-Seq study (Ulrich et al. 2022). In addition, we generated in-house and utilized additional publicly available scRNA-Seq data from adjacent non-malignant tissues and tumors from patients with HGSCs (total: n = 88 samples across 58 donors, **Fig 1A, Suppl. Table 1**). We further clustered 9,759 stromal cells, 32,598 T/NK cells, and 1,850 myeloid cells from 12 FT samples (Dinh, Lin, et al. 2021) (**Fig 1B**, **Suppl. Fig 1A-B**) and identified 7 myeloid, 4 T cell, 2 NK cell, 4 fibroblast, 2 pericyte, and 1 smooth muscle cell subsets (**Methods**). In addition, using label transfer (Stuart et al. 2019), we were able to identify all annotated cell subsets in an independent scRNA-Seq dataset from 4 benign FT donors (Ulrich et al. 2022) (**Suppl. Fig 1C-D**). Our analysis identifies greater myeloid diversity measured by an entropy-based scoring method, Rogue (Liu et al. 2020) compared to T/NK cell clusters despite being just 5% as frequent as T/NK cells in our scRNA-Seq (**Suppl. Fig 1E**) samples. Additionally, we calculated the number of unique statistically differentially expressed (DE) genes **(Suppl. Fig 1F**), showing that myeloid cells had consistently more DE genes per cluster and justified the decision to maintain more myeloid clusters compared to T/NK cells. In our stromal subsets, we identify 3 fibroblast subsets and a small cluster with a myeloid-like gene signature, with potential antigen-presenting functions as recently identified in multiple tissue types (Elyada et al. 2019; Dinh, Pan, et al. 2021). Due to the limited number of captured cells in the scRNA-Seq, we did not further cluster other cell types, such as B and endothelial cells.

**Figure 1.**
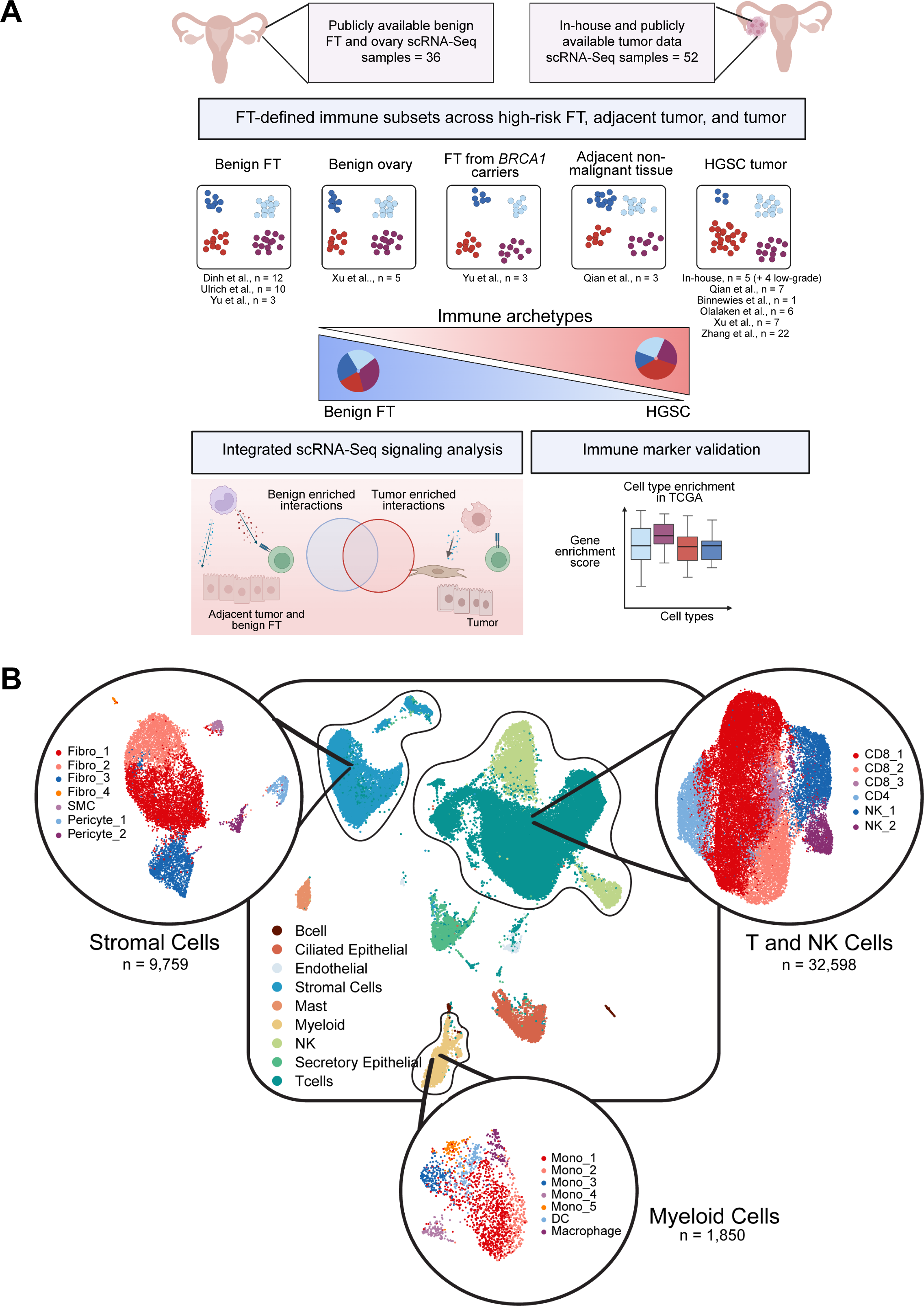
Integrated scRNA-Seq analysis of human fallopian tubes to identify immune features of HGSC progression. A. Bioinformatics analysis of 88 scRNA-Seq samples, including in-house and publicly available data, identifies heterogeneity and interactions of immune and stromal cells in FTs and their alterations across adjacent normal tumor, benign ovary, and HGSCs. (Created with Biorender.com) B. Transcriptional heterogeneity of stromal, T/NK, and myeloid cell subsets from 12 FT samples (Dinh, et al. 2021) served as a reference for further analysis across relevant tissue and sample types.

### Myeloid cell heterogeneity and plasticity in human FTs

Myeloid cell diversity in high-grade serous ovarian cancers was recently described with multifaceted, functional gene signatures in distinct tumor microenvironments (TME) (Hornburg et al. 2021). We reasoned that an in-depth transcriptional characterization of FT myeloid cells would help us identify a link between the myeloid microenvironment of HGSC and FTs, the tissue of origin for most HGSC tumors. Our clustering analysis (**Methods**) defined 4 classical CD14+ monocyte subsets, 1 nonclassical CD16+ monocyte subset, 1 dendritic cell, and 1 macrophage subset (**Fig 2A**) that were shared across FT samples (**Suppl. Fig 1B**). Downstream analysis including differential gene expression, gene set enrichment analysis using JASMINE (Noureen et al. 2022), and gene regulatory network analysis from SCENIC (Aibar et al. 2017) (**Methods**) defined distinct gene signatures (**Fig 2B-D**) in the 7 myeloid subsets and supported our cell type annotations. We used representative markers to annotate myeloid subsets, including 4 classical monocyte subsets with variable expression levels of *CCL4*, a chemoattractant of T and myeloid cells that was found to correlate with CD8+ and FOXP3+ T cell infiltration in HGSCs <u>(Zsiros et al. 2015)</u>. Other markers include the proteoglycan versican (*VCAN*) and heat shock proteins (HSPs) describing stress-responsive monocytes (Mujal et al. 2022). Macrophages, which expressed complement complex genes *C1QA/B* as well as *TREM2* were transcriptionally distinct from monocytes by their lack of *FCN1* and *S100A8/9* expression as well as chemoattractant and inflammatory molecules such as *IL1B*, *CXCL2/3/8*, and *CCL20*. JASMINE pathway analysis using Gene Ontology Biological Process (GO: BP) revealed gene sets enriched for classical monocytes (inflammatory response, leukocyte chemotaxis), HSP+ monocytes (protein folding), non-classical CD16+ monocytes (interferon-inducible pathways), antigen-presenting pathways in macrophages and DCs and lipid metabolism pathways unique to macrophages (**Fig 2C**). Gene regulatory network analysis performed with SCENIC <u>(Aibar et al. 2017; Van de Sande et al. 2020)</u> showed *BHLHE41* and *MAF* regulons enriched within macrophages; both negative regulators of macrophage proliferation and activation (Aziz et al. 2009; Rauschmeier et al. 2019). *EHF* and *ETV3* regulons were also active, likely reflecting dendritic cell differentiation programs (Appel et al. 2006; Villar et al. 2023), and *IRF4*, a DC-lineage transcription factor that promotes TH2 polarization (Williams et al. 2013) (**Fig 2D**).

**Figure 2.**
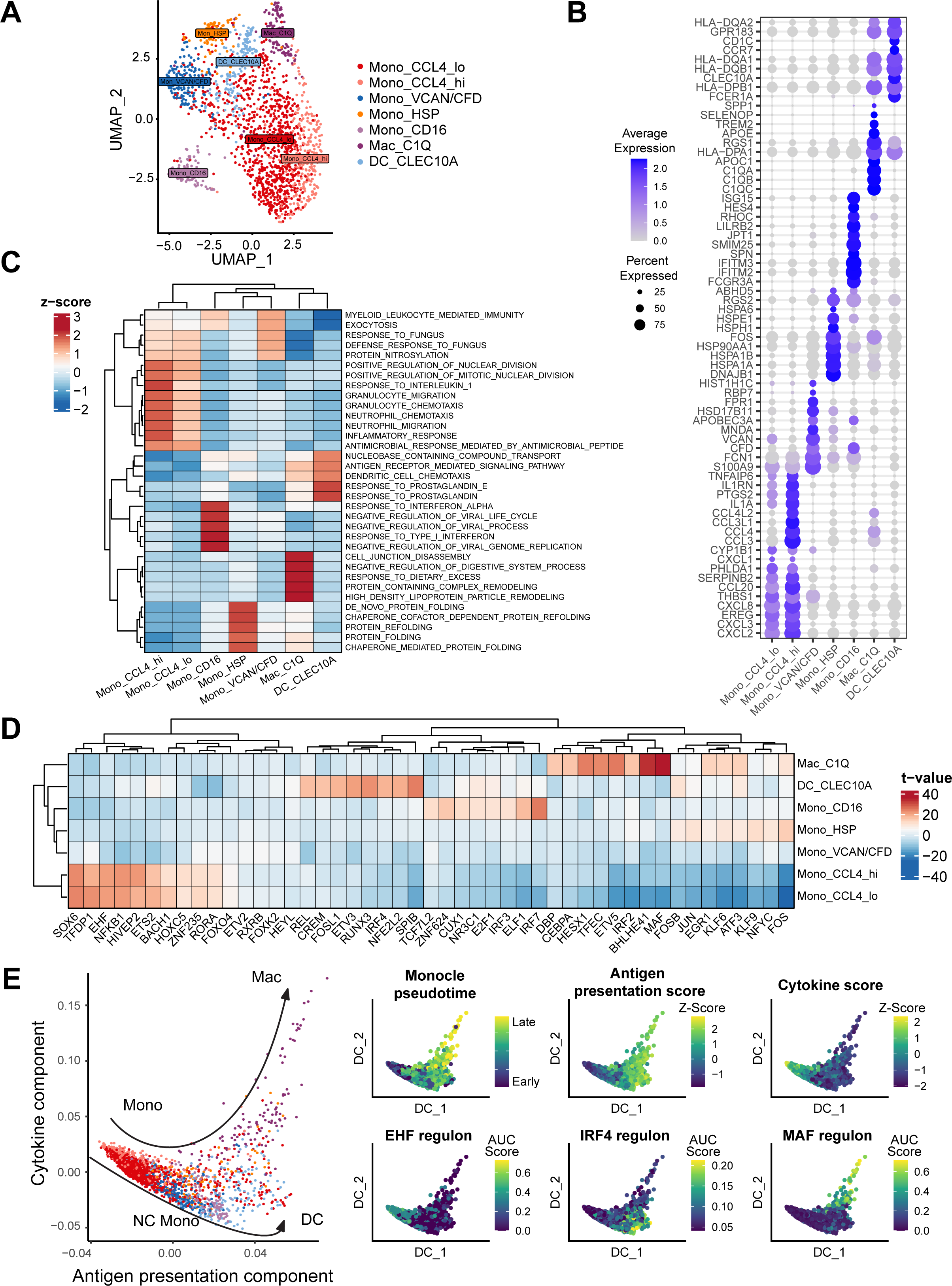
Myeloid cell heterogeneity in human FTs shows higher diversity of monocytes compared to macrophages and dendritic cells. A. UMAP and clustering of integrated myeloid cells identify 4 classical monocyte subsets (*CD14*, *FCGR3A*-), 1 subset of non-classical monocytes (*CD14*, *FCGR3A+*), a subset of type 2 dendritic cells (*CD1C, CLEC10A*), and macrophages (*APOE*, *TREM2*). B. Representative gene signatures of myeloid subsets presented by DotPlot; values show scaled RNA expression across clusters. C. Gene set enrichment analysis of top scoring Biological Processes Gene Ontology (GO: BP) for myeloid subsets. Values shown are z-scores of Jasmine’s odds-ratio test across clusters. D. Gene regulatory network analysis using SCENIC shows regulon scores enriched for each myeloid subset. Values plotted are derived from linear models using one cluster vs. the remaining and extracting the t-values for each model (Methods). E. Diffusion map of myeloid cells calculated using the top 2000 variable features. Monocle3 pseudotime, gene set scores, and SCENIC AUC scores were overlaid on the 1^st^ two diffusion components.

To test how potential monocytic developmental trajectories in human FT correlates with our defined regulatory networks, we paired SCENIC results with diffusion maps <u>(Angerer et al. 2016)</u> and Monocle’s graph-based pseudotime <u>(Cao et al. 2019)</u>. Diffusion map analysis suggested two distinct diffusion components, one associated with MHC-II antigen presenting genes and the second with inflammatory cytokine expression **(Fig. 2E**, **Suppl. Fig 2A**). Monocle supported the inferred pseudotime trajectory from classical CD14+ monocytes to more matured monocytic cell types (dendritic cells, macrophages) or distinct monocyte states (HSP/*VCAN* monocytes and non-classical CD16+ monocytes) supported by the enrichment score of SCENIC regulons. The top correlated genes with each diffusion component indicated how monocytic cells might function differently during their maturation. We ranked the top 10 correlating genes with each diffusion component (**Suppl. Fig 2A**) to indicate how monocytic cells might function distinctly during their maturation. For instance, enrichment scores derived from the average expression in single-cell clusters suggested cytokine/interleukin genes were enriched with monocytes while HLA gene sets were enriched within macrophages and DCs.

Next, we used FT-defined myeloid cell signatures to evaluate their enrichment in 394 HGSC samples from TCGA (Cancer Genome Atlas Research Network 2011) and found that signatures of more mature monocytic subsets such as *VCAN*+ monocytes and macrophages were most enriched for immunoreactive and mesenchymal HGSC subtypes (**Suppl. Fig 2B, Methods**). Together, our analysis revealed extensive heterogeneity of myeloid cells in human FTs and suggested a phenotypic change in myeloid cells that may be relevant in HGSCs.

### Macrophage-to-monocyte ratio is distinct in the FT, adjacent normal tissues, and HGSC tumors

Previous flow cytometry and scRNA-Seq analyses did not annotate monocyte subset in FTs (Ardighieri et al. 2014; Ulrich et al. 2022); however their signatures were identified in HGSC scRNA-Seq (Hornburg et al. 2021) that correlated with T/NK cell phenotypes. Since monocyte signatures (*S100A8/9*, *FCN1*, *VCAN*) from circulating blood (Villani et al. 2017) and HGSC tumor (Hornburg et al. 2021) are consistent with our FT-defined monocytes, we asked if we could identify monocytes in independent samples of healthy FT scRNA-Seq (Ulrich et al. 2022) (**Suppl. Fig 3A- B**). Our clustering analysis recovers abundant monocytes with similar marker genes as identified in our FT samples (*S100A8/9*, *FCN1*, *TREM2*, *CD1C*) and consistently differentiated macrophages, monocytes, and DCs from each other. Using label transfer (Stuart et al. 2019), we show that monocyte subsets are retained in other previously published data (Ulrich et al. 2022) (**Suppl. Fig 3C-D**). We observed that *S100A9* and *FCN1* consistently separate monocytes from dendritic cells and *TREM2*+ *C1QA/B* macrophages. The comparatively low abundance of macrophages relative to monocytes in benign FT scRNA-Seq datasets led us to evaluate their proportions in adjacent normal tissues compared to HGSC tumors from the same patients. We reanalyzed two independent scRNA-Seq datasets, one with paired adjacent normal and tumor from <u>(Qian et al. 2020)</u> and another with 5 benign ovarian tissues and 7 HGSCs from <u>(Xu et al. 2022).</u> We found a significantly elevated macrophage-to-monocyte ratio in HGSC tumors compared to benign FT tissues (3.40 fold increase, FDR adjusted p-value = 3.6×10^-6^) (**Fig 3A- B**), while the difference between adjacent, non-malignant tissue, or benign ovary compared to healthy FT tissue are less pronounced (1.90, and 1.94 fold increase, respectively) and were not found to be statistically significant (FDR adjusted p-values = 0.19, 0.50). The macrophage-to-monocyte ratio was not significantly different between adjacent normal tissues compared to that in low-grade ovarian cancer in our newly generated scRNA-Seq (**Fig 3B**). Taking advantage of the recent availability of published scRNA-Seq datasets, we reanalyzed additional HGSC tumor scRNA-Seq <u>(Xu et al. 2022; Binnewies et al. 2021; Olalekan et al. 2021)</u> and, on average, found a higher macrophage-to-monocyte ratio in HGSC TME from different datasets (total n = 26 HGSC samples).

**Figure 3.**
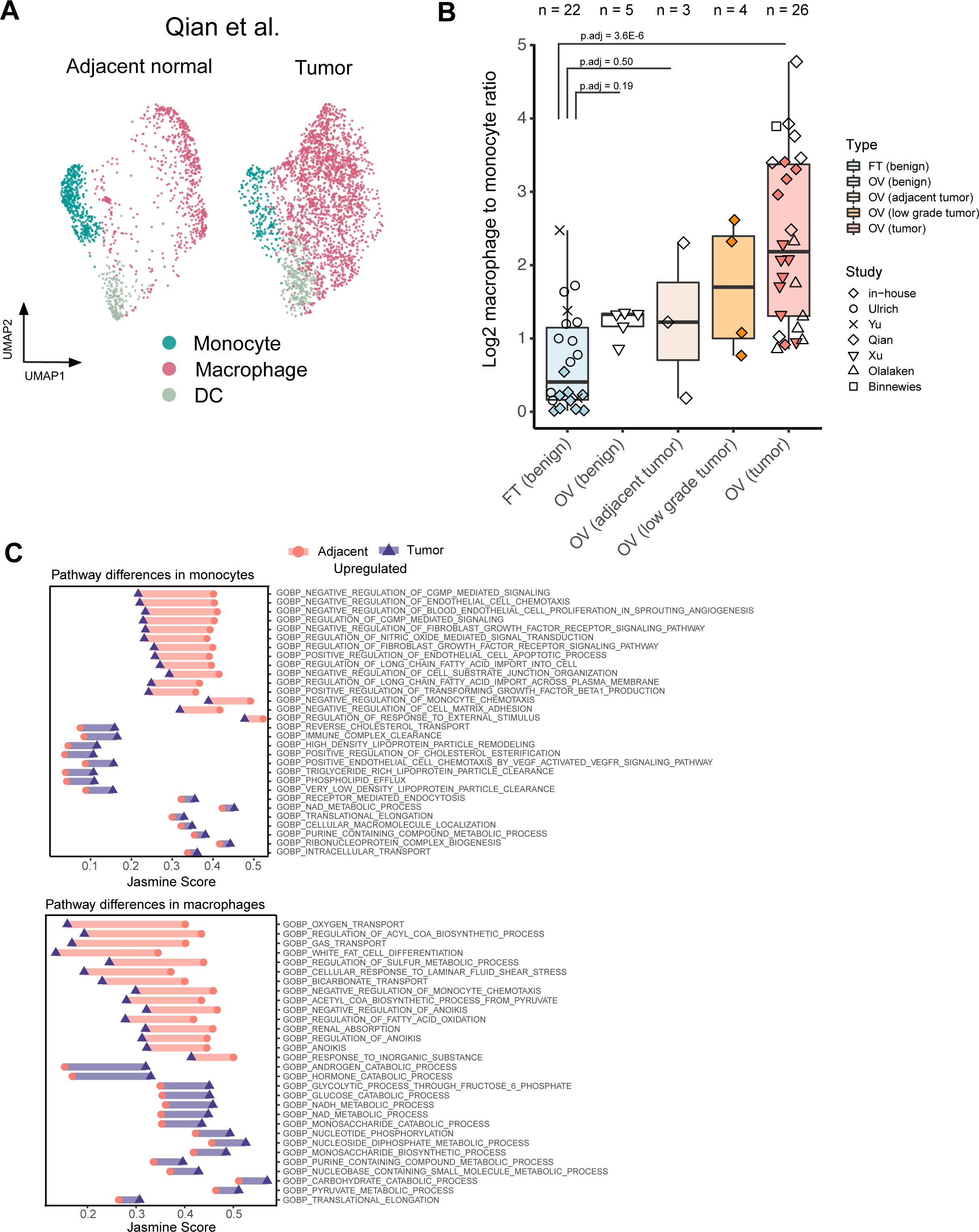
Analysis of macrophage-to-monocyte ratio across tissue types and conditions. A. UMAP and clustering of myeloid cells from publicly available data (Qian et al. 2020) shows the enrichment of macrophages relative to monocytes. (n = 3 samples per group, 2 patients). B. The log2 macrophage-to-monocyte ratio across benign FT, benign ovary, adjacent normal tumor, low-grade, and HGSC tumor (FDR-adjusted p-value, two-sided pairwise Wilcox test). C. Jasmine scores (GO:BP) calculated for macrophages and monocytes across tumor adjacent and tumor microenvironments from Qian et al. scRNA-Seq (panel A). Significant pathways (two-sided t.test, FDR-adjusted p-value < 0.01) were ranked by fold change between microenvironments and the Jasmine scores plotted.

To identify the potential difference in myeloid cells’ functions in adjacent normal and benign tissue, we used scRNA-Seq of adjacent normal (n = 3) and HGSC tumors (n = 7) (Qian et al. 2020). JASMINE GO: BP pathway analysis showed marked differences in monocytes in adjacent tissues compared to tumors, including enrichment of anti-angiogenic pathways in adjacent monocytes compared to tumors (negative regulation of endothelial cell chemotaxis FDR adjusted p-value = 1.58 ×10^-25^ and positive regulation of endothelial cell apoptotic process FDR adjusted p-value = 4.10×10^-21^) (**Fig 3D**). In contrast, tumor-associated monocytes were significantly enriched in metabolic-related pathways (High-density lipoprotein particle remodeling FDR adjusted p-value = 4.20 x10^-14^ and NAD metabolic process FDR adjusted p-value = 1.33 x10^-9^) compared to adjacent normal monocytes, and similar to tumor-associated macrophages (TAMs). We also observed shifted metabolic pathways in TAMs, supporting their change in energy demands compared to adjacent tissue. Notably, we see a reduced reliance on oxygen for metabolism demonstrated with a significantly lower JASMINE score for the ‘oxygen transport’ pathway (FDR adjusted p-value = 5.11 x10^-265^) and ’acetyl coenzyme-A biosynthesis’ pathway (FDR adjusted p-value = 9.03×10^-211^), which is an essential precursor for oxidative phosphorylation.

### FT-defined cell-cell interactions of monocytes and macrophages are correlated in adjacent normal and matched HGSC tumors

Since the phenotypes of monocytes and macrophages are associated with different types of HGSC TMEs <u>(Hornburg et al. 2021)</u>, we asked if the cell-cell interactions of monocytes and macrophages with other cell types in FTs are altered in tumor compared to benign tissue (FT and adjacent normal) (**Fig 3B**). To test this, we further identified 6 T/NK cell subsets, including 3 CD8+ cell subsets, 2 NK cell subsets, and 1 CD4 subset with distinct gene signatures (**Suppl. Fig 4A- B**). Among those, scRNA-Seq-derived signature scores identified T/NK subsets were highest in immunoreactive subtypes in most subsets, but particularly in NK cells. (**Suppl. Fig 4C**). Further sub-clustering did not obtain more subsets with sufficient DE genes for T cell subset annotation with conventional phenotypic markers, including T memory and helper subtypes (**Suppl. Fig 4D**).

**Figure 4.**
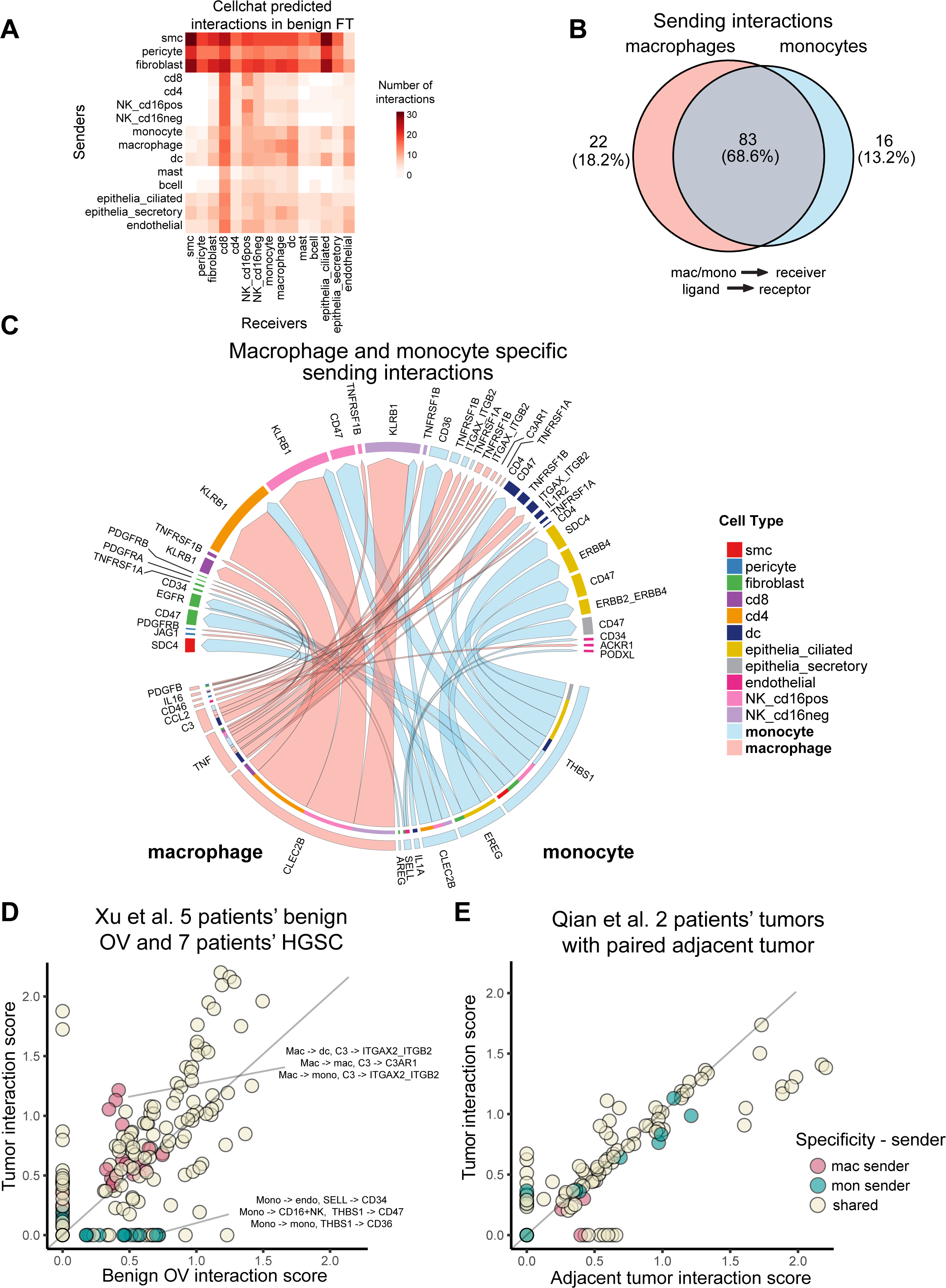
Cell-cell interaction analysis of monocytes and macrophages in FTs with other cell types. A. Frequency of significant CCIs in human FTs (12-samples, Dinh et al. 2021) identified by CellChat’s permutation test. B. Comparison of monocyte- and macrophage-specific sending CCIs in human FTs. C. Circos plot of monocyte- and macrophage-specific CCIs. Arrow width represents Cellchat’s scaled interaction scores. D. CCI coexpression scores of monocyte- and macrophage-specific CCIs in tumor and benign ovary (Xu et al. 2022). Each point represents a CCI score. E. CCI coexpression scores of monocyte- and macrophage-specific CCIs in in tumor and adjacent tumor samples (Qian et al. 2020).

To evaluate the cell-cell interaction (CCI) signals from scRNA-Seq, we used Cellchat (Jin et al. 2021). ligand-receptor database and quantified the significant interactions across major subtypes in our defined human FT atlas (**Fig 4A**). CellChat’s permutation test identified 121 significant ligand-receptor pairs with monocytes/macrophages as a sender cell type, in which 83 (68.6%) were shared between macrophages and monocytes, while 22 (18.2%) and 16 (13.2%) specifically identified from either macrophages or monocytes, respectively (**Fig 4B-C**). The dominant signaling pathways of macrophage CCIs were *CLEC2B* and TNF signaling, which engages the *KLRB1* and TNFR receptors, known for self-recognition and negative regulation of T/NK cell activation (Iizuka et al. 2003; Mathewson et al. 2021) and inflammation, respectively. Other macrophage CCIs, including *PDGFB-PDGFRA/B* between macrophages and stromal cells, were reported in HGSC tumors with reactive stroma and poor prognosis (Li et al. 2022). On the other hand, monocyte CCIs included epiregulin (*EREG*), which has potential interactions with stromal cells through *EGFR*, or ciliated epithelial cells through HER family receptors *ERBB2* and *ERBB4*, reported in low-grade serous ovarian cancers (Luo et al. 2018). Another relevant monocyte interaction, *THBS1-CD47*, showed a reduction in angiogenesis and increased tumor rejection in a xenotransplant model, suggesting its pro-tumoral effects (Jeanne et al. 2021). Next, we performed the same CCI analysis using macrophages and monocytes as receiving populations (**Suppl. Fig 4E-F**), which highlighted the increased diversity in macrophage receptor use such as with *LRP1*, *AXL*, *CSF1R*, *HAVCR2*, *C3AR1* when compared to monocytes.

Since we observed the difference in the macrophage-to-monocyte ratio in samples compared to tumors, we asked if the FT-defined CCIs expressed differently in adjacent normal and tumors. To do so, we compared co-expression CCI scores (**Methods**) of the FT-defined monocyte- and macrophage-specific and shared CCIs in adjacent normal and tumor samples from two independent scRNA-Seq datasets (Xu et al. 2022; Qian et al. 2020). The co-expression of our CCI scores positively correlated with *CellChat*’s communication probabilities (**Suppl. Fig 4G)**, which allowed us to identify cross-tissue CCIs between benign, adjacent normal, and tumor tissues. We observed that interactions with monocytes as senders were enriched in the benign ovary (Xu et al. 2022), with 11 monocyte-specific positive scored interactions in benign ovary and 2 in tumors, whereas most interactions with macrophages as senders were similar between normal and tumor (**Fig 4D**). In a second data set we observed a high correlation between these interactions, but not in a cell type-specific manner (Qian et al. 2020). The receiving signals were also consistently correlated between these datasets (**Suppl. Fig 4H-I**), indicating that tumor microenvironment and cell-cell interactions may be intrinsic to their cell identities based on their high correlation across different tissues.

### Monocyte and macrophage diversity in patients with germline *BRCA1* mutations and HGSC patients treated with chemotherapy

We identified the macrophage-to-monocyte ratio changes potentially linked with disease progression. Next, we wanted to evaluate this ratio using FT scRNA-Seq from patients with *BRCA1* germline mutations (Yu et al. 2022), who are high risk of developing HGSCs. We used marker genes consistent with our 12 sample FT reference to identify myeloid cell types (**Suppl. Fig 5A-B**). We observed an increase in macrophage-to-monocyte ratio in FT of *BRCA1* germline mutation (p-value = 0.7, two-sided Wilcox test, n = 3 vs 3). (**Fig 5A**). Differential gene expression showed an increased in the chemokines *CCL3* and *CXCL2/8*, first identified from our FT monocyte subsets, in macrophages and monocytes of BRCA1 FT samples, and decreased expression of heat shock protein (HSP) genes in BRCA samples (**Fig 5B**). The data indicated a potential of change in macrophage-to-monocyte ratio in FT of HGSC high-risk patients compared to healthy FT but remains limited due to small sample size.

**Figure 5.**
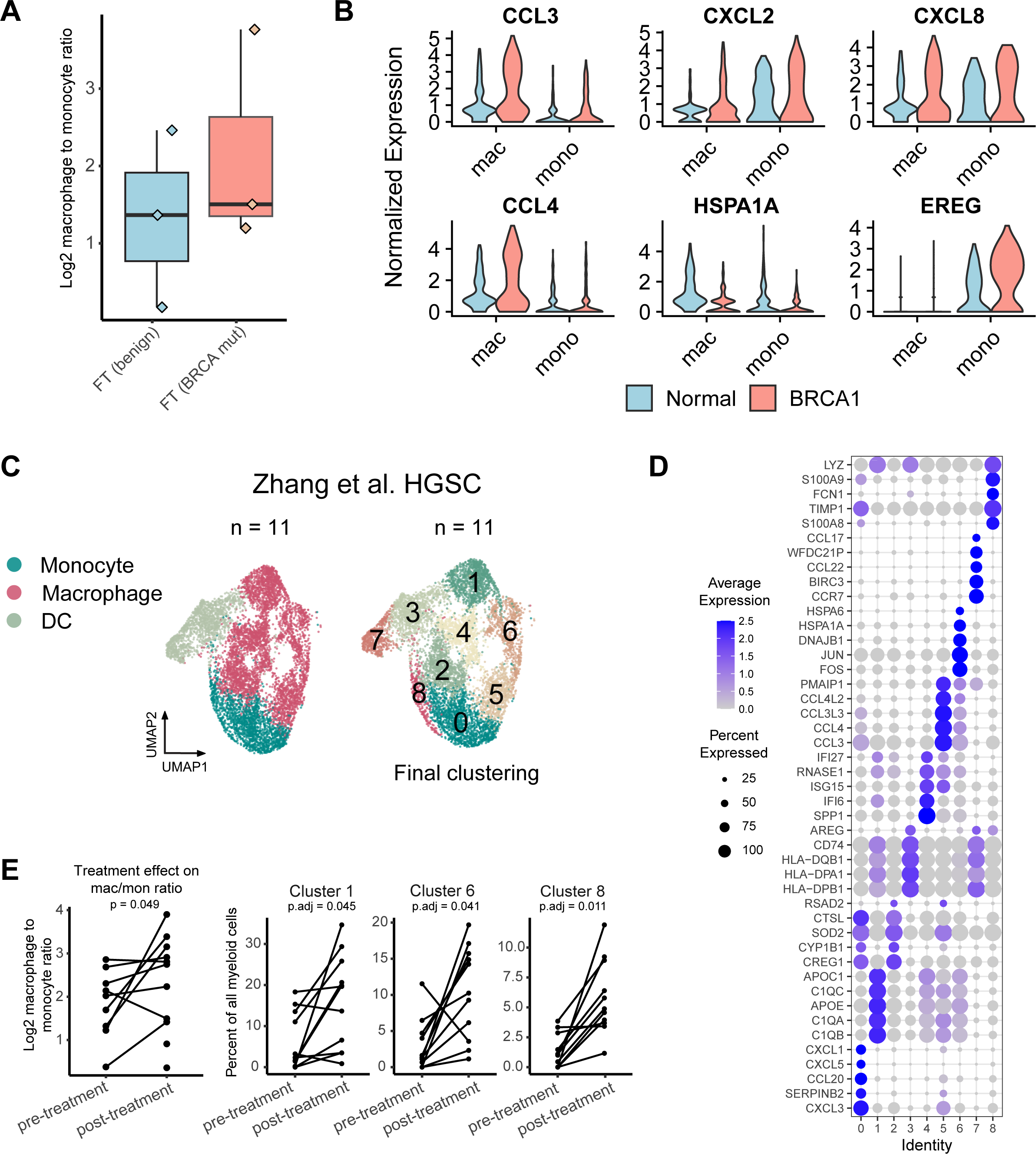
Macrophage and monocyte heterogeneity analysis shows the altered gene expression in high-risk patients with germline BRCA mutations and response to chemotherapy. A. The log2 macrophage-to-monocyte ratio in benign FT compared with FT in germlin*e BRCA1* carriers (data from Yu et al. 2022.). B. Examples of monocytic genes defined in human FTs are differentially expressed in FTs of *BRCA*1 carriers. Values shown are normalized RNA counts. C. UMAP and clustering reanalysis of myeloid cell subsets from 11 donors pre- and post-neoadjuvant chemotherapy (Zhang et al. 2022). D. Differential expression analysis of myeloid subsets, values shown are z-scored expression across clusters. E. Evaluation of macrophage-to-monocyte ratios, and abundance of myeloid subsets from paired pre-and post-chemotherapy treated samples. Clusters 1 and 6 (macrophages) and cluster 8 (monocytes) were significantly enriched post-treatment (two-sided paired t.test, FDR-adjusted p-values < 0.05).

We were also curious how macrophage-to-monocyte ratio may change in HGSC patients upon treatment. We reanalyzed a publicly available treatment-naïve scRNA-Seq dataset from 11 patients taken before and after neoadjuvant chemotherapy treatment (Zhang et al. 2022) (**Suppl. Fig 5C**). Our myeloid cell clustering identified 2 populations of monocytes, 5 populations of macrophages, and 2 dendritic cell subsets (**Fig 5C-D**) with distinct gene signatures resembling those defined in our FT analysis (*CCL3/4*, *CXCL3*, *HSPA1*, *FCN1*, *CCL20*, *C1QA*). Overall, chemotherapy’s effect on the macrophage-to-monocyte ratio is uncertain. There is no significance when using all samples, including those with poor cell recovery, which are likely more variable (p-value = 0.456). However, after removing samples with fewer than 50 total myeloid cells combined (n = 2), we see a 1.45-fold increase in the log2 ratio (p-value = 0.0485). More samples are needed to determine the reproducibility of this effect. Additionally, we observed three subsets that significantly increased in proportion after treatment, including macrophages expressing *APOE*, *APOC1*, and complement components (C1Qs) (cluster 1, FDR adjusted p-value = 0.045), HSP+ stress-related macrophages (cluster 6, FDR adjusted p-value = 0.041), and classical monocytes (cluster 8, FDR adjusted p-value = 0.011), that were most similar to the early FT-defined monocyte subsets (**Fig 5E**). Several subsets of monocyte and macrophage subpopulations are changed in HGSC as expected **(Suppl. Fig 5D**), despite the uncertain impact on the ratio of macrophage-to- monocytes. We also note the increased complexity of myeloid cell diversity in the TME, especially within macrophages relative to benign tissue.

### Stromal-immune cell interactions in human FTs

Since stromal cell-cell interactions were predicted to be the most frequent sending cell types inferred in FT scRNA-Seq, (**Fig 4A**), we asked to what extent their interactions with immune cells differ between normal FTs and HGSC tumors. To do so, we sub-clustered the stromal cell compartment and identified three fibroblast subsets, two pericyte populations, and one smooth muscle cell (SMC) subset (**Fig 6A**) with distinct gene expression signatures, including complement genes (*C7*, *C3*, *CFD*) and cytokine/chemokine expression (*CXL8*, *IL6*) (**Fig 6B**). We named the subsets with representative markers except for SMCs, which was comprised of one subset. F_RASD1 fibroblasts were suspected to be less differentiated/activated with high regulon activity of stemness markers *SOX4* and *TWIST2* from SCENIC (**Suppl. Fig 6A)** (Yang et al. 2020). Smooth muscle and pericyte populations could be characterized by differential expression of cell adhesion molecules (*MCAM*), *RGS5* (pericyte markers), *MYH11*, and *ZCCHC12* as smooth muscle cell markers **(Suppl. Fig 6B).** Among those subsets, we found that F_C7 fibroblasts, marked by high expression of complement *C7* and SMC signature scores, were enriched in the TCGA HGSC mesenchymal subtype, known to have the worst survival outcomes (**Fig 6C)**. Next, we used cancer-associated fibroblast (CAF) signatures from HGSC scRNA-Seq (Hornburg et al. 2021; Olbrecht et al. 2021) to evaluate FT stromal subsets. We find complement expression (*CFD*) and other genes (*DPT*, *MGP*, and *CXCL12*) similar to inflammatory fibroblasts described in HGSCs (Hornburg et al. 2021) were enriched in our F_C7 subset relative to other stromal cells. (**Suppl. Fig 6C**). CCI analysis using CellChat identified interactions between stromal subsets and immune receiver cells, particularly through *CD44*, *CD47* and integrins (**Suppl. Fig 6D**). Most CCIs from fibroblast to immune cells were shared between our three fibroblast subsets (80%), though F_C7 fibroblasts have 22 CCIs not shared with other fibroblast subsets (12% of all stromal to immune interactions) (**Fig 6D**). Notably, fibroblast to myeloid signaling, including *C3* to *C3AR1,* associated with immunosuppression and cancer cell proliferation (Huang, Zhou, and Deng 2023), and *C3* to *ITGAX* and *ITGB2*, were among these (**Suppl. Fig 5D**). *FN1* interactions, which were defined from our C7 expressing fibroblasts show interactions with diverse immune cell subsets and were predicted to have higher ligand-receptor co-expression in tumor relative to benign tissue from 2 independent cohorts (**Fig 5E-F**). Altogether, we reported a diversity of stromal cell subsets in human FTs, with distinct gene expression profiles and subsets that may contribute to distinct interactions enriched in HGSCs.

**Figure 6.**
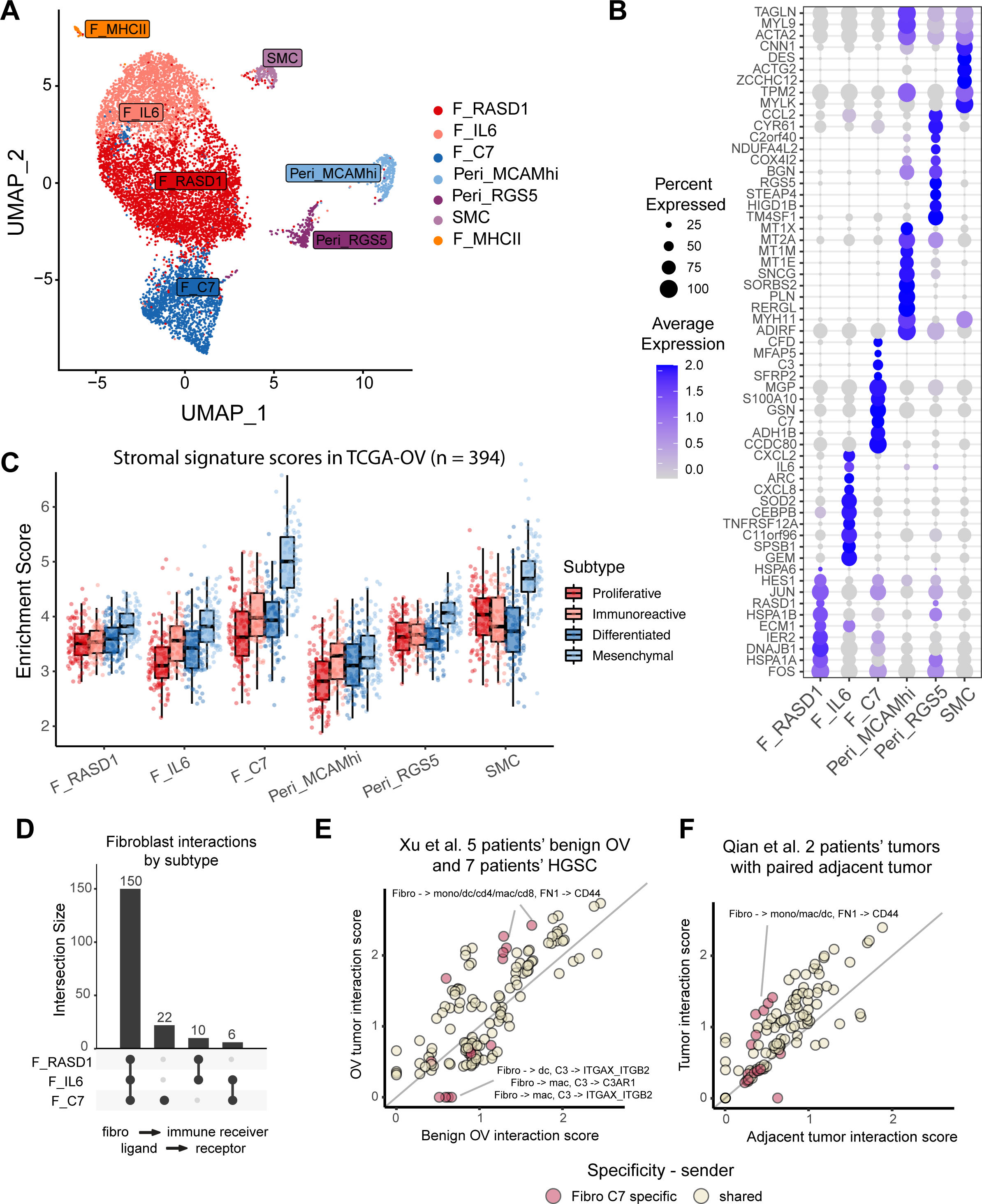
Stromal heterogeneity analysis of human FTs shows enrichment of complement-expressing fibroblasts and smooth muscle cells in mesenchymal HGSC subtype. A. UMAP projection of 3 fibroblast subsets, 1 smooth muscle cell subset, and 2 pericyte subsets. B. Top 10 DE genes of stromal subsets presented by DotPlot; expression values are z-scaled across clusters. C. Stromal subset signature enrichment analysis on TCGA-OV samples (n=394). D. Upset plot shows overlapping ligand-receptor interaction numbers from fibroblast subsets to immune cells, values derived from Cellchat’s permutation test. E. Stromal CCI co-expression analysis in HGSCs vs. benign ovary (Xu et al. 2022). F. Stromal CCI co-expression analysis in HGSCs vs. tumor adjacent (Qian et al. 2020).

## Discussion

This study characterizes the diversity of non-epithelial compartments in benign FTs, especially immune cells and cell-cell interactions which was not fully described from recent scRNA-Seq. The defined single-cell transcriptional landscape of FT immune cells showed a greater extent of heterogeneity compared to the previous characterization using flow cytometry (Ardighieri et al. 2014). An independent, human scRNA-Seq dataset (Ulrich et al. 2022) shows the cell types identified here were reproducible. Specifically, we identified 5 fibroblast and smooth muscle cell subsets and 2 pericyte subsets, including a rare subset expressing antigen-presenting markers recently found in several tissues and cancer types. A fibroblast subset expressing *C7* was transcriptionally similar to an inflammatory CAF signature found in scRNA-Seq of HGSCs (Hornburg et al. 2021) and was most enriched in mesenchymal subtypes of TCGA compared to our other stromal subsets. CCI analysis predicted stromal cells as having the most frequent interactions in human FT with unique ligand-receptor interactions defined in *C7* fibroblasts. These interactions included fibronectin (*FN1*) which correlates with reactive stroma and recurrence after chemotherapy treatment (Ryner et al. 2015). The *FN1* interactions (*FN1*-*CD44*) from fibroblasts to immune cells were also increased in tumors relative to non-malignant tissues.

We identified high monocyte diversity in FT with similar gene expression profiles to those observed in circulation (*S100A8/9*, *FCN1*, *VCAN*, *CD14*) (Mulder et al. 2021; Villani et al. 2017). Five monocyte subsets were identified along with DCs and macrophages with distinct transcriptional signatures of chemokines, antigen-presentation, transcription factor usage (SCENIC), and pseudotime trajectories. We provided a comprehensive analysis of monocyte heterogeneity which has yet to be reported in human FT tissues, highlighting the importance of single-cell profiling technology. When compared to benign tissue, we observed a significant increase in the macrophage-to-monocyte ratio in HGSC tumors. Furthermore, monocytes and macrophages have specific ligand-receptor interactions with other cell types, along with many shared interactions defined by scRNA-Seq. Monocytes with *EREG* expression were predicted to interact with epithelial expressing *ERBB2/4*, a growth factor that might play a role in tumor progression, and macrophages expressing *PDGF*, associated with aggressive HGSC stroma (Li et al. 2022), that may interact with fibroblasts could indicate a role in cancer-associated fibroblast formation in the tumor microenvironment. Interestingly, monocyte-specific interactions are found more frequently in benign tissues, while sone macrophage-specific were higher in HGSC tumors in recent scRNA-Seq data of 12 patients (Xu et al. 2022). However, most ligand-receptor pairs defined in monocyte and macrophage interactions were positively correlated in benign and tumor tissues and in another scRNA-Seq dataset of 2 patients with matched tissues (Qian et al. 2020). Further work is needed to evaluate the mechanisms of monocytic maturation and recruitment in HGSC progression. We suspect that early changes in tumor development may differentiate monocytes in a tissue-specific manner. Others have shown that the co-culture of PBMC-derived monocytes with ovarian cancer cell lines was sufficient for alternative macrophage development through TGF-alpha (Fogg et al. 2020). We hypothesize that the macrophage-to-monocyte ratio could be an indicator of HGSC progression. This concept was recently applied within a melanoma and renal cell carcinoma context (Mujal et al. 2022), showing that the macrophage-to-monocyte ratio positively correlated with tumor Treg infiltration. Recently published scRNA-Seq of HGSCs also showed a correlation between *FCN1* expressing monocytes and “immunological desert” phenotypes compared to HGSCs with T-cell infiltrated or excluded tumors (Hornburg et al. 2021). Lastly, we evaluated monocyte and macrophage heterogeneity in two publicly available datasets: scRNA-Seq of 3 women with germline BRCA1 mutations with 3 age-matched samples (Yu et al. 2022) and 11 HGSC patients with pre and post-neoadjuvant samples (Zhang et al. 2022). We observed the trend of macrophage-to-monocyte ratio in BRCA1 samples, but the low sample size (n = 3 per condition) limited this analysis. However, we found that BRCA1 samples had higher pro-inflammatory markers, including chemokine expression in their monocytes (*CCL3/4*, *CXCL2/8*) and *EREG* signaling, also found in our CCI analysis. Similarly, only when excluding low abundant samples in the chemotherapy response data (Zhang et al. 2022), we observed a statistically significant change in the macrophage-to-monocyte ratio between pre- and post-treatment samples, with 2 macrophage and 1 classical CD14+ monocyte subsets that were increased after treatment. This data indicates the important but less explored roles of monocytes and macrophages in HGSC progression and response to treatment.

With poor survival outcomes and a lack of early markers for detection, we consider the possibility of immune or stromal signatures as candidates for early disease progression. This work contributes to characterizing the phenotypic changes of non-epithelial cell types in human FT and their impacts on HGSC development that promote further studies that lead to immune-based biomarkers of early cancer. We defined the extent of monocyte diversity in FTs, previously overlooked in tissue as they’re typically considered circulating cell types. Emerging data has supported their correlation with tumor phenotypes and potential functions (Mujal et al. 2022; Mulder et al. 2021; Cheng et al. 2021). The observed transcriptomic differences of monocytes and macrophages in adjacent and tumor tissue and the change in their proportion support their importance in HGSC and potentially other tumors.

Our work has several limitations. The lack of clinical data, such as survival from publicly available HGSC scRNA-Seq data, limits the correlation analysis of immune and stromal cell diversity inferred from FT scRNA-Seq. The limitation of other data modalities, like high-dimensional flow cytometry, prevents the current findings from further exploration and validation. Additionally, the ability to power these analyses and independently validate the gene expression profiles of the diverse cell subsets was aided by publicly available data; however, we note some the limitations to this approach. We analyzed these datasets independently as we expect cell type diversity to be lost through a computational dataset integration (CCA, harmony, or similar). We expect dataset-specific biases to be present and cannot verify how differences in sample preparation may have influenced the recovery of cell type proportions. Additionally, multiple samples were derived from different regions within a single patient but were treated as independent in our framework. In total, we had 58 unique patients and 88 total scRNA-Seq samples. The calculation of cell type ratios will be inherently noisy relative to other cell types (stromal cells, T cells), as myeloid cells were less abundant. Finally, others have also shown a significant correlation of CD68+ cells in normal fallopian tube epithelia during the luteal phase (George, Milea, and Shaw 2012), and lack the clinical metadata of our samples to identify and account for this trend.

## Materials and Methods

### Biological sample handling and processing for scRNA-Seq

In-house scRNA-Seq data was generated from high-grade or low-grade serous ovarian tumor patients with informed consent and approval of the Institutional Review Board at Cedars-Sinai Medical Center (**Suppl. Table 1**). Fresh human primary tissues were placed in sterile serum-free MEM at 4 °C and transferred to a tissue culture laboratory. Tissues were minced into ∼1–2 mm pieces before digesting with 1 × Collagenase/Hyaluronidase (STEMCELL Technologies, Catalog #: 07912) and 100 μg/mL DNase I (Sigma Aldrich, SKU: 10104159001) in 7 mL of serum-free MEM. The samples were incubated at 37 °C with constant rotation for 90 min. The supernatants were collected and the cell suspensions were spun down at 300 g for 10 min at 4 °C. To lyse red blood cells, the cell pellets were resuspended in red blood cell lysis buffer (0.8% NH_4_Cl, 0.1% KHCO_3_, pH7.2) and incubated for 10 min at room temperature. Cell suspensions were spun again at 300 g for 10 min at 4 °C and the cell pellets were resuspended in PBS. If >5% dead cells were observed by trypan blue staining, dead cell removal was performed using dead cell removal kit (Miltenyi Biotec, Catalog #: 130-090-101) according to the manufacturer’s instructions. Remaining cells were directly used for single cell RNA sequencing (scRNA-Seq) or frozen in 90% fetal bovine serum supplied with 10% dimethyl sulfoxide in a Mr. Frosty freezing container placed at −80 °C. Frozen cell vials were transferred to gas phase of liquid nitrogen for long-term storage. Cells were thawed and transferred into a 15 mL conical tube with 7 mL of serum-free medium and then spun down at 300 g for 10 min at 4 °C. The cell pellets were resuspended in 100 µL of PBS. Cells were counted using a hemocytometer and the sample volume was adjusted to achieve a cell concentration within 100-2,000/µL.

### Single-cell capture, library preparation, and next-generation sequencing

Single cells were captured and barcoded using the 10X Chromium platform (10X Genomics). scRNA-seq libraries were prepared following the instructions from the Chromium Single Cell 3ʹ Reagent Kits User Guide (v3). Briefly, Gel Bead-In EMulsions (GEMs) were generated using single-cell preparations. After GEM-RT and cleanup, the complementary DNAs (cDNAs) from barcoded single-cell RNAs were amplified before quantification using Agilent Bioanalyzer High Sensitivity DNA chips. The single-cell 3′ gene expression libraries were constructed and cDNA corresponding to an insertion size of ∼ 350-400 bp were selected. Libraries were quantified using Agilent Bioanalyzer High Sensitivity DNA chips and pooled together to get similar numbers of reads from each single cell before sequencing on the NovaSeq S4 (Novogene).

### ScRNA-Seq pre-processing of publicly available data and in-house HGSC samples

We downloaded single-cell RNA-seq data from publicly available sources through the NCBI GEO, Zenodo, or like resources (**Suppl. Table 1**). Depending on the origin of the source (either gene count matrix or processed data in the Seurat objects), we reanalyzed publicly available data using count matrices for consistent processing. We performed QC and filtered out the low-quality cells with >15% mitochondrial UMIs or fewer than 800 transcripts. Then, we used the single-cell transform (SCTransform) method (Hafemeister and Satija 2019) for expression normalization and regressed out the percent mitochondrial reads and cell cycle scores. Principal components analysis (PCA) was performed from the “SCT” assay prior to integration analysis to remove unwanted technical variance using the Harmony method <u>(Korsunsky et al. 2019)</u> with default settings. The corrected PCA (Harmony components) were used to generate UMAP embeddings and clusters.

### Single-cell integration analysis of benign FT samples

Non-epithelial compartments (stromal, myeloid, and T/NK cells) were subset from our previously published dataset (Dinh, Lin, et al. 2021) to generate new Seurat objects. Each cell subset was used further integrated analysis across different donors using default 2000 features Integration anchors were calculated using “SCT” as normalization method. The *k.filter* parameter was manually set for myeloid cells and fibroblasts to maintain more donors, but kept to default in T/NK subsets where cells were most abundant. Four samples were excluded for myeloid cell analysis due to the low number of captured cells (**Suppl. Table 1**).

To validate the presence of cell subsets, we used Label Transfer from Seurat to identify our annotated cluster in an independent cohort of benign FT samples (Ulrich et al. 2022). The two datasets were normalized using SCT and the top 3000 most variable genes were used for PCA. Predicted annotations using the *FindTransferAnchors* and *TransferData* functions. To quantify the similarity between two datasets, the top 3000 variable features were used for pairwise correlation analysis of our clusters with the predicted cluster labels and plotted using the Pheatmap package (Kolde, R. 2012.).

### Single-cell clustering, markers, and subset identification

We identified cell clusters using Seurat graph-based clustering methods at increasing resolutions from 0.1 to 1 to identify major cell types (T/NK, stromal, B, myeloid cells) within a scRNA-Seq dataset. We used marker genes - *LYZ*, *CD14*, *CD68*, *FCN1*, *CLEC10A*, *CD1C* - to identify myeloid cells, which were subset and converted into a new Seurat object for further clustering. We re-performed normalization and integration as described on the subset data to identify macrophages and monocytes. Macrophages were characterized by *C1Q*A/B/C, *TREM2*, *APOE*, and *HLA-DRA* high expression and monocytes by *FCN1*, *S100A8/9*, and *FCGR3A* for non-classical monocytes. We required a cluster with a minimum of 5% percent of cells from the total cell population. We use the Clustree package <u>(Zappia and Oshlack 2018)</u> to visualize and guide splitting or merging clusters with the following criteria: a minimum of 5 differentially expressed genes using MAST (Finak et al. 2015), with a Bonferroni-corrected p-value < 0.01, log2FC > 0.25, and a percent expression difference greater than 25% from the new cluster and its parent in previous clustering resolutions. If these criteria were not met at a clustering resolution, we would merge the cluster back into its parent population and leave the remaining clusters unchanged.

### Quantifying heterogeneity of myeloid and T/NK cells with differential gene expression and entropy

We used the *Rogue* method (Liu et al. 2020) to quantify the proportion of genes with significant entropy from all expressed genes in a scRNA-Seq sample. We used the RNA counts for myeloid and T/NK cell subsets for entropy scoring and performed initial filtering to require a minimum of 10 cells expressing a gene to be considered for the entropy scoring. Entropy for all donors per cell cluster were calculated using the *SE_fun* function. Rogue scores represent the number of genes determined to have a significant entropy divided by the total number of genes expressed and ranges from 0 to 1. Rogue values were not estimated for samples with fewer than 10 cells per cluster.

In addition, we used DE analysis (adjusted p-value < 0.01, gene expressed in >20% of cells) for all resolutions from 0.1 to 1 using *FindAllMarkers* function in Seurat package. At that given clustering resolution, we sum the unique DE genes of the given subset (myeloid, T/NK, or stromal cells) while excluding mitochondrial and ribosomal genes (MT-, RPS, RPL), divided by the number of unique clusters at that clustering resolution.

### ScRNA-Seq signature enrichment analysis in TCGA data

Using the top 10 DE genes per cluster from scRNA-Seq, we computed the average expression (Log normalized counts of transcript per million expression) in TCGA bulk RNA-Seq samples. Two-sided pairwise Wilcox tests were computed for each cluster across the 4 molecular subtypes, using Benjamini-Hochberg FDR p-value correction for multiple comparisons.

### JASMINE pathway analysis

Gene set enrichment analysis was performed using Jasmine’s odds ratio test <u>(Noureen et al.</u> <u>2022)</u>. The input data took in the RNA count matrices and used GO:BP genesets from the Gene Ontology database through *MsigDB* (https://www.gsea-msigdb.org/gsea/msigdb). This method was robust against dropouts relative to other gene set scoring methods. We ranked genesets by significance using general linear models and linear contrasts to identify gene sets that were most significant for a given cluster compared to the mean of other clusters. We selected the top 5 pathways per cluster by highest t-value, then z-scaled Jasmine enrichment scores for heatmap visualization after filtering out non-relevant pathways. Enrichment score comparisons were performed using t-test with FDR p-value correction using the Benjamini-Hochberg method. We ranked the pathways by the absolute change by average Jasmine score.

### Gene regulatory network analysis (SCENIC)

We applied SCENIC (Aibar et al. 2017) to infer gene regulatory networks in stromal, T/NK, and myeloid cells in benign FT. We used the human hg38 reference with 10kb upstream and downstream from the transcription start sites to search for DNA motifs. AUC scores generated for each regulon were used to create linear models with linear factors with the Multcomp package (v1.4-16; Hothorn et al., 2008). for hypothesis testing. Linear contrasts were constructed using cluster against the mean of the remaining clusters and selected those with the highest t-values. We selected the top regulons for each cell cluster and extracted the T values, representing the cluster-specific enrichment for each regulon that and plotted using ComplexHeatmap (Gu Z, 2022) in R.

### Pseudotime/trajectory inference analysis

We used the diffusion map method in the “Destiny” R package <u>(Angerer et al. 2016)</u>, from log-normalized RNA counts from the top variable genes. Spearman’s correlation test identified genes associated with the two diffusion map dimensions representing the potential cellular trajectories. We also used *Monocle3* (Cao et al. 2019) to infer the pseudotime of single cells to independently support the gene expression patterns observed in diffusion map analysis.

Antigen presentation scores were calculated by taking the averaged log expression of all HLA genes, and CD74 expressed in a minimum of 10% of cells from any given myeloid cell cluster. Similarly, cytokine scores were calculated using cytokine genes (CXCL-, CCL-, IL-) expressed in 10% of any myeloid cluster. The scores were then overlaid in the inferred trajectories.

### Cell-cell communication and co-expression ligand-receptor score

To identify potentially interacting cell types, we applied the CellChat method <u>(Jin et al. 2021) to</u> <u>predict ligand-receptor interactions with</u> default parameters (FDR adjusted p-value < 0.05). We filtered out interactions with CD4 as receptors (with MHCII) on myeloid cell types. We used the predicted macrophage- and monocyte-specific interactions to quantify how ligand-receptor pairs may be co-expressed in different tissues using a similar scoring scheme using the geometric mean from robust means (½ median + ¼*(first + third quartiles)) of each ligand and receptor expression for each cell cluster.

### Macrophage-to-monocyte ratio across datasets

The macrophage-to-monocyte ratio was calculated by the sum of macrophage clusters divided by monocytes per sample in each dataset. We add 1 as a pseudo-count and log2-transform to prevent negative values. Significance was tested using a two-sided Wilcox rank test and corrected using the Benjamini-Hochberg method. In datasets with paired pre- and post-treatment (Zhang et al. 2022), we used a two-sided paired t.test.

### Data & Code availability

All source code generated in this study has been uploaded to a Zenodo repository and will be made available at time of peer-reviewd publication. Raw FASTQ files and h5 files (count matrices) of in-house scRNA-Seq data has been uploaded to NCBI GEO and will be made available at time of peer-reviewed publication.

## Acknowledgements

This project was partially supported by the RIDE scholar award 2021 (H.Q.D). The research in the Dinh lab was supported by startup packages from the Human Cancer Genetics and Precision Medicine cluster, Carbone Cancer Center, UW School of Medicine and Public Health, Vice Chancellor for Research and Graduate Education and Center for Human Genomics and Precision Medicine. Research reported in this publication was supported by the National Institute Of General Medical Sciences of the National Institutes of Health under Award Number T32GM135119 (J.B). The content is solely the responsibility of the authors and does not necessarily represent the official views of the National Institutes of Health.

---

**Suppl. Figure 1.**
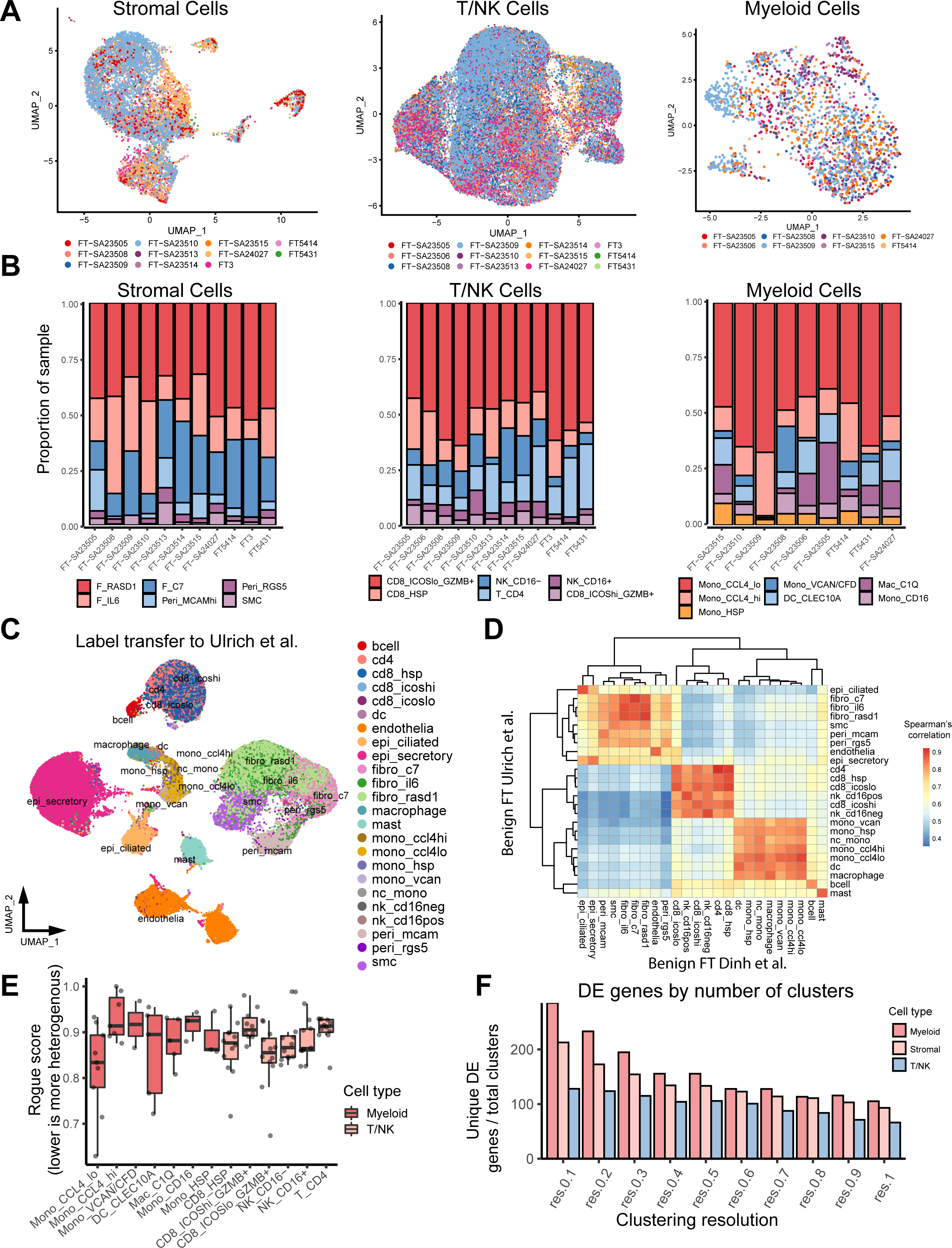
A. Sample overlay on UMAPs of CCA-integrated scRNA-seq of benign FT samples for stromal, T/NK, and myeloid cell compartments. B. Proportions of cell clusters across donors in stromal, T/NK, and myeloid cell compartments. C. Reclustering and label transfer predictions of Ulrich et al. scRNA-Seq data based on 12- FT-sample reference using SCT normalization and harmony integration. D. Pairwise correlation of the top variable genes (n = 2000) from FT reference and the predicted cell annotations in benign FT (Ulrich et al. 2022). E. Entropy-based ‘Rogue’ scores measure the heterogeneity of individual scRNA-seq FT samples across clusters. F. Differential expression calculated at increasing clustering resolutions for myeloid, stromal, and T/NK cells. The number of unique differentially expressed genes (adjusted p-values < 0.01) calculated with MAST, were normalized by the number of clusters per clustering resolution.

**Suppl. Figure 2.**
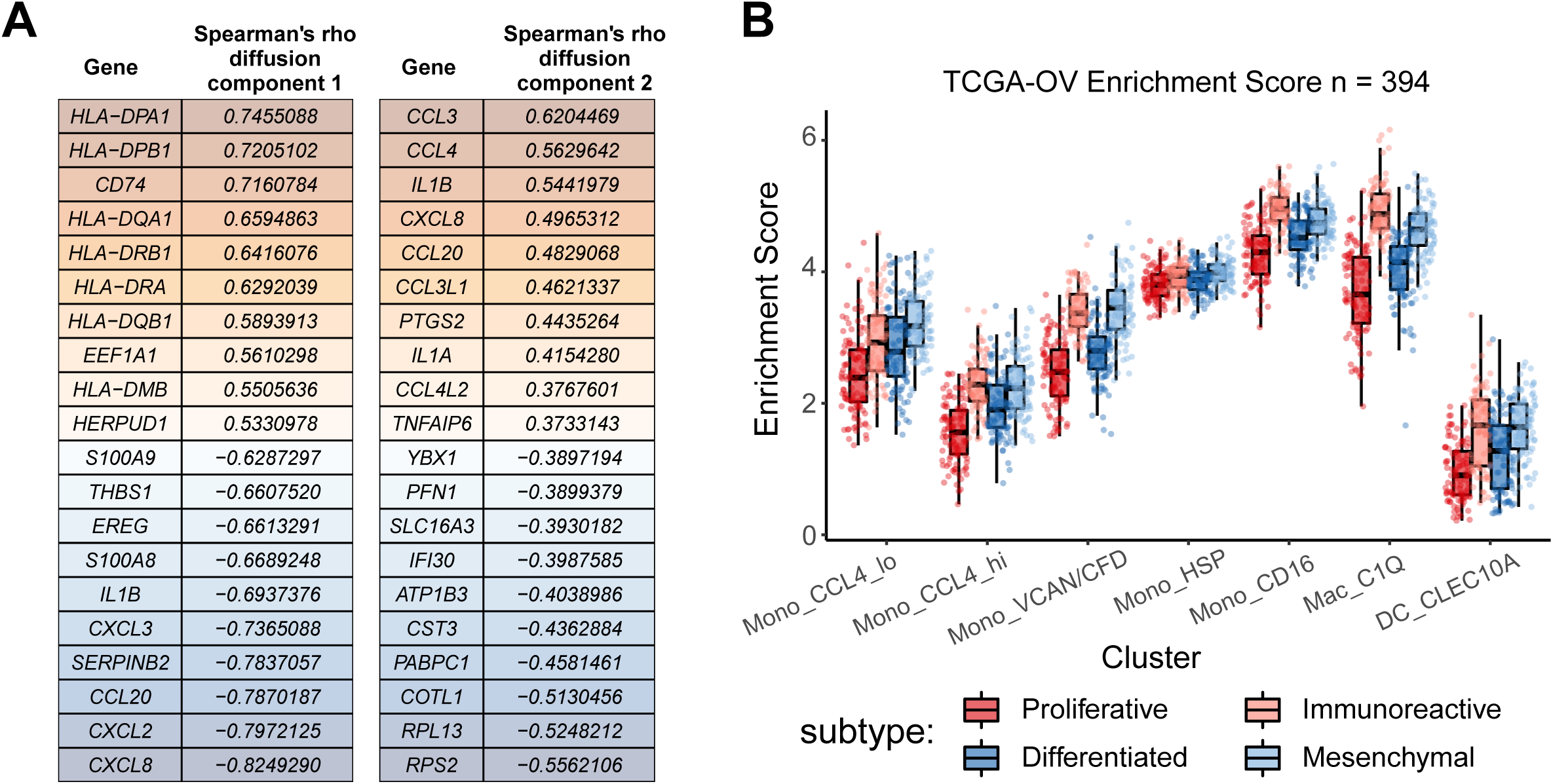
A. Spearman’s rho of the top 10 positively and negatively correlated genes for the first 2 diffusion components show enrichment for cytokine and antigen presentation genes. B. ScRNA-Seq gene signature analysis of bulk RNA sequencing samples from 4 molecular subtypes of TCGA HGSC samples (n=394) (Methods).

**Suppl. Figure 3.**
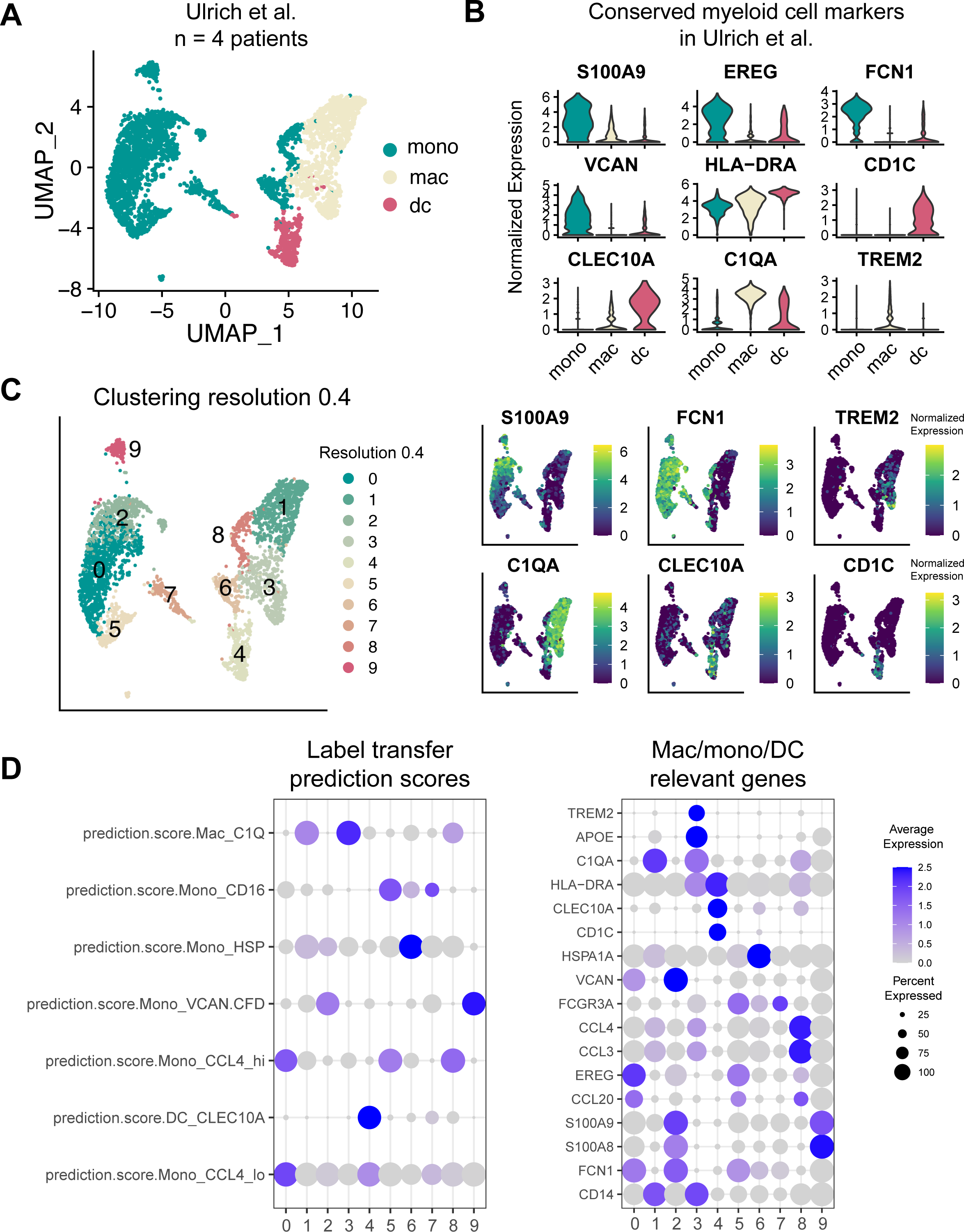
A. Reclustering and annotating myeloid cells from benign FT from 4 cancer-free patients (Ulrich et al. 2022). B. Myeloid gene signatures support gene expression specificity between 12 FT sample reference data and data from Ulrich et al. for identifying macrophages, monocytes, and DCs. C. Further clustering of myeloid cells identifies heterogeneity and guided cell type annotation, normalized gene expression shown. D. Label transfer predictions for myeloid clusters in independent data (Ulrich et al., 2022) using 12 sample FT reference. Gene signatures of the predicted clusters presented in a DotPlot, with z-scaled expression values.

**Suppl. Figure 4.**
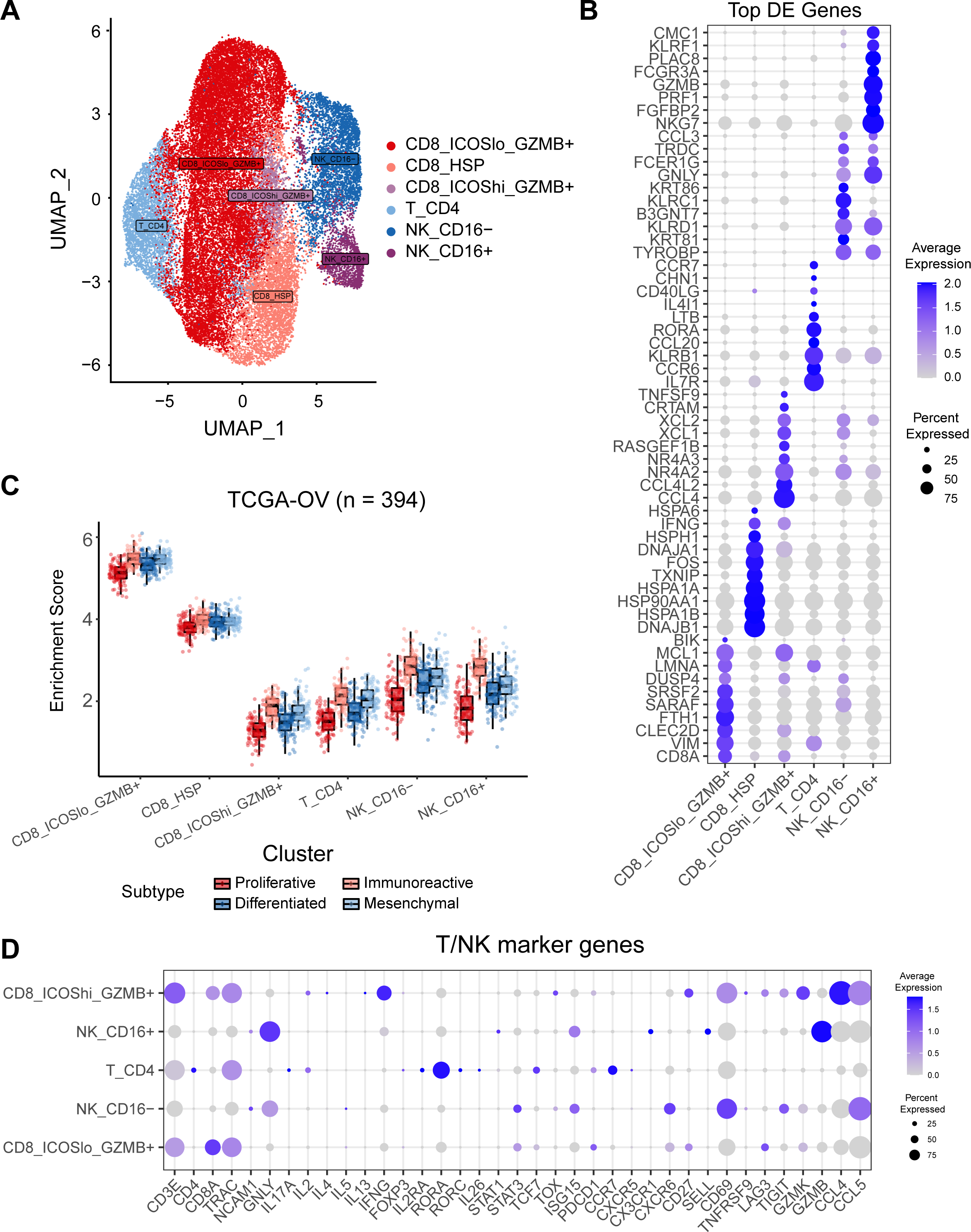
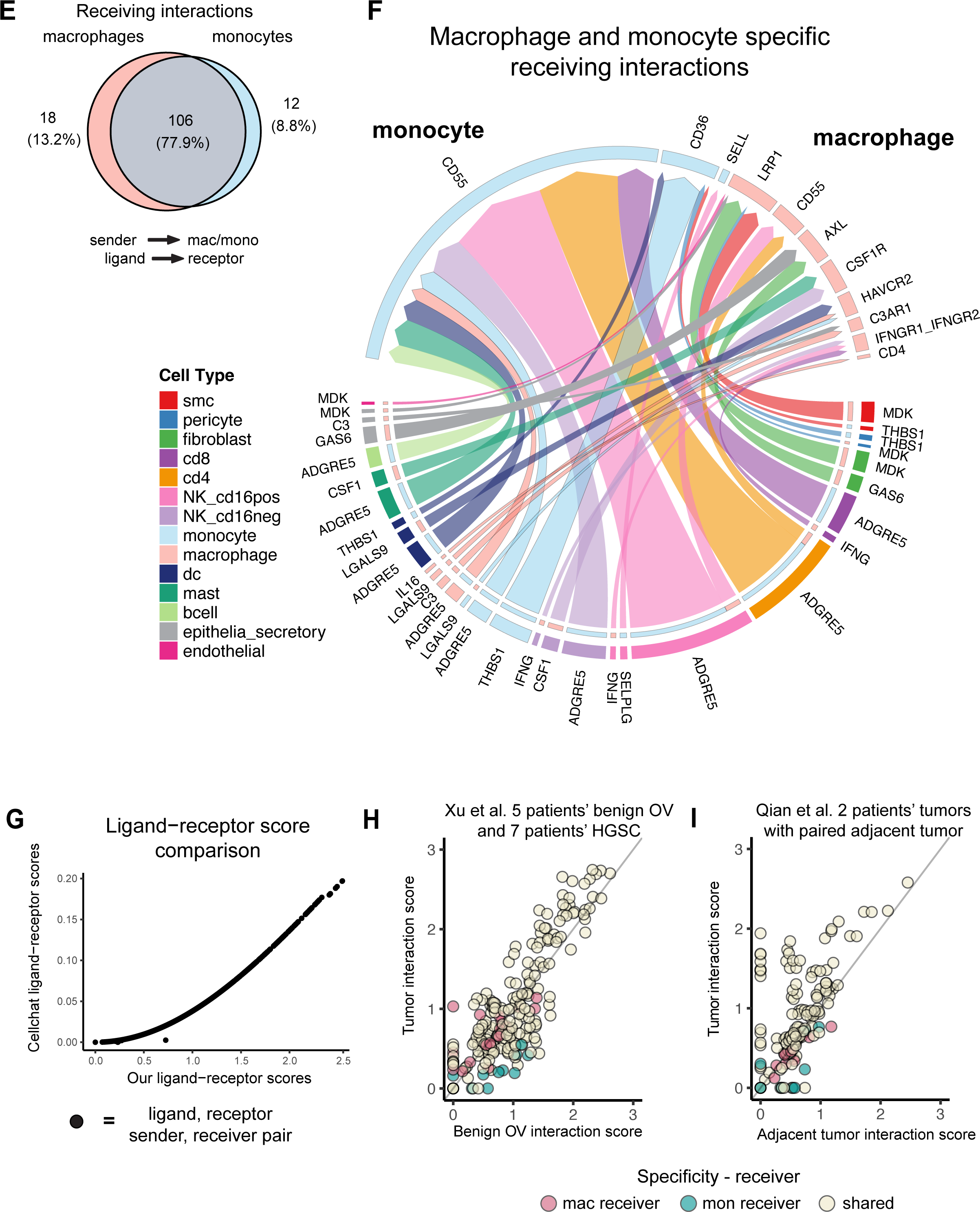
A. Clustering of T/NK cells identifies 3 CD8, 1 CD4, and 2 NK cell subsets projected on the UMAP. B. Gene signatures for T/NK cell clusters presented by DotPlot, values shown are z-scaled expression. C. Signature enrichment analysis of T/NK cell subsets for TCGA-OV samples (n=394). D. T cell memory and activation gene signatures in FT T/NK cell subsets. E. CCI with macrophages and monocytes as receiver cells in benign FT. F. Circos plot presentation of monocyte- and macrophage-specific interactions (as receivers). G. Comparison of CellChat CCI score and co-expression score. Values shown are from each possible ligand-receptor, sender-receiver pair using Cellchat’s human interaction database. H. Macrophage and monocyte receiving CCIs across benign ovary, HGSC, and adjacent tumor (Xu et al. 2022). I. Macrophage and monocyte receiving CCIs across adjacent tumor and HGSC tumors (Qian et al. 2020).

**Suppl. Figure 5.**
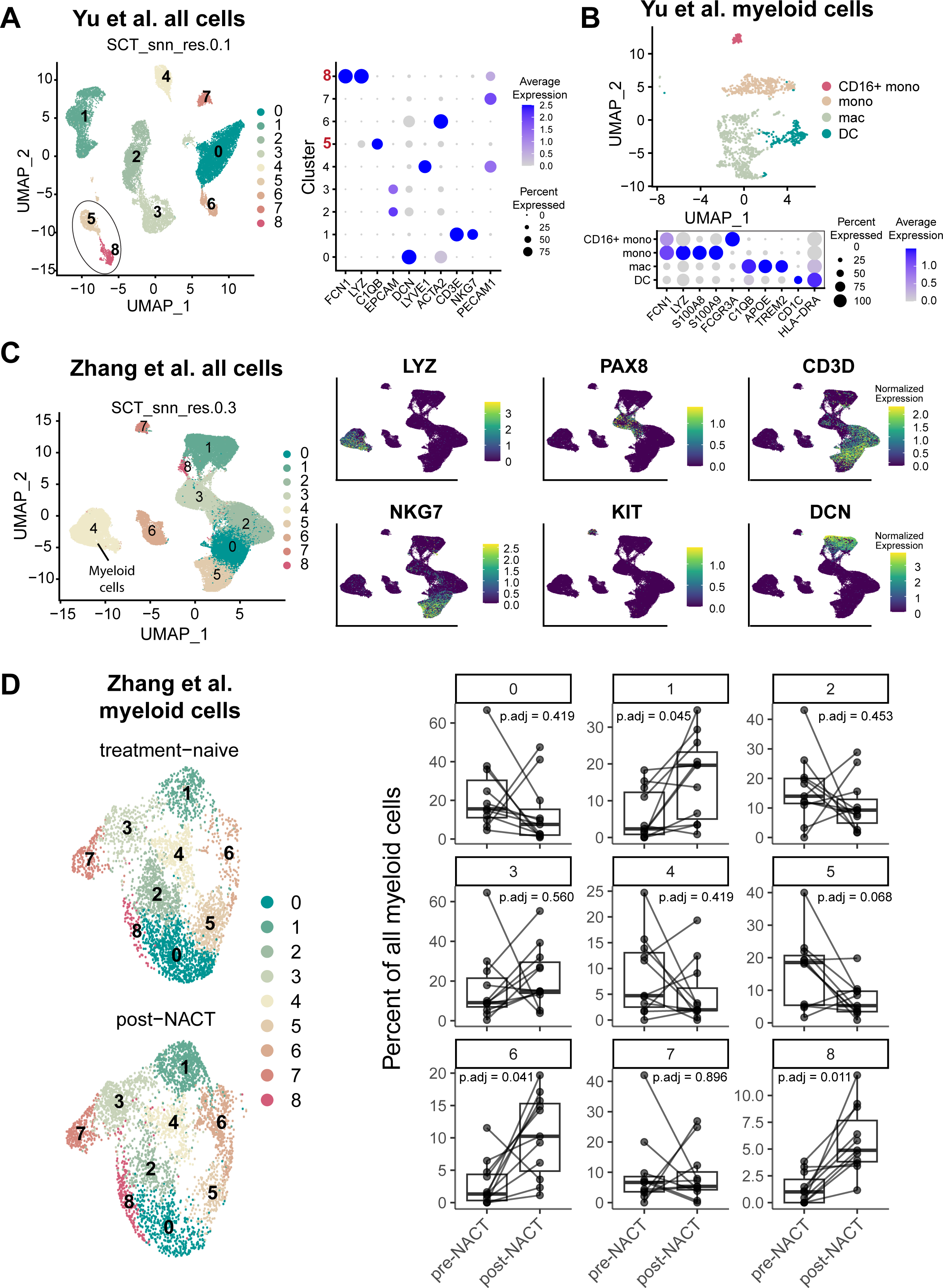
A. ScRNA-Seq analysis of myeloid cells, clusters 5 and 8 expressing *LYZ*, *FCN1*, and *C1QB,* were selected for further clustering analysis. A. B. Gene signatures of myeloid subsets (z-scored expression) identifies DCs, macrophages, non-classical, and classical monocytes. B. Integrated analysis of pre- and post-chemotherapy treated patients (Zhang et al. 2022). Myeloid cells (*LYZ*+) were selected for further clustering analysis. C. UMAP projection of myeloid cells for pre- and post-treatment HGSC patients and comparison of cluster frequencies (two-sided paired t.test, FDR-adjusted p-values shown).

**Suppl. Figure 6.**
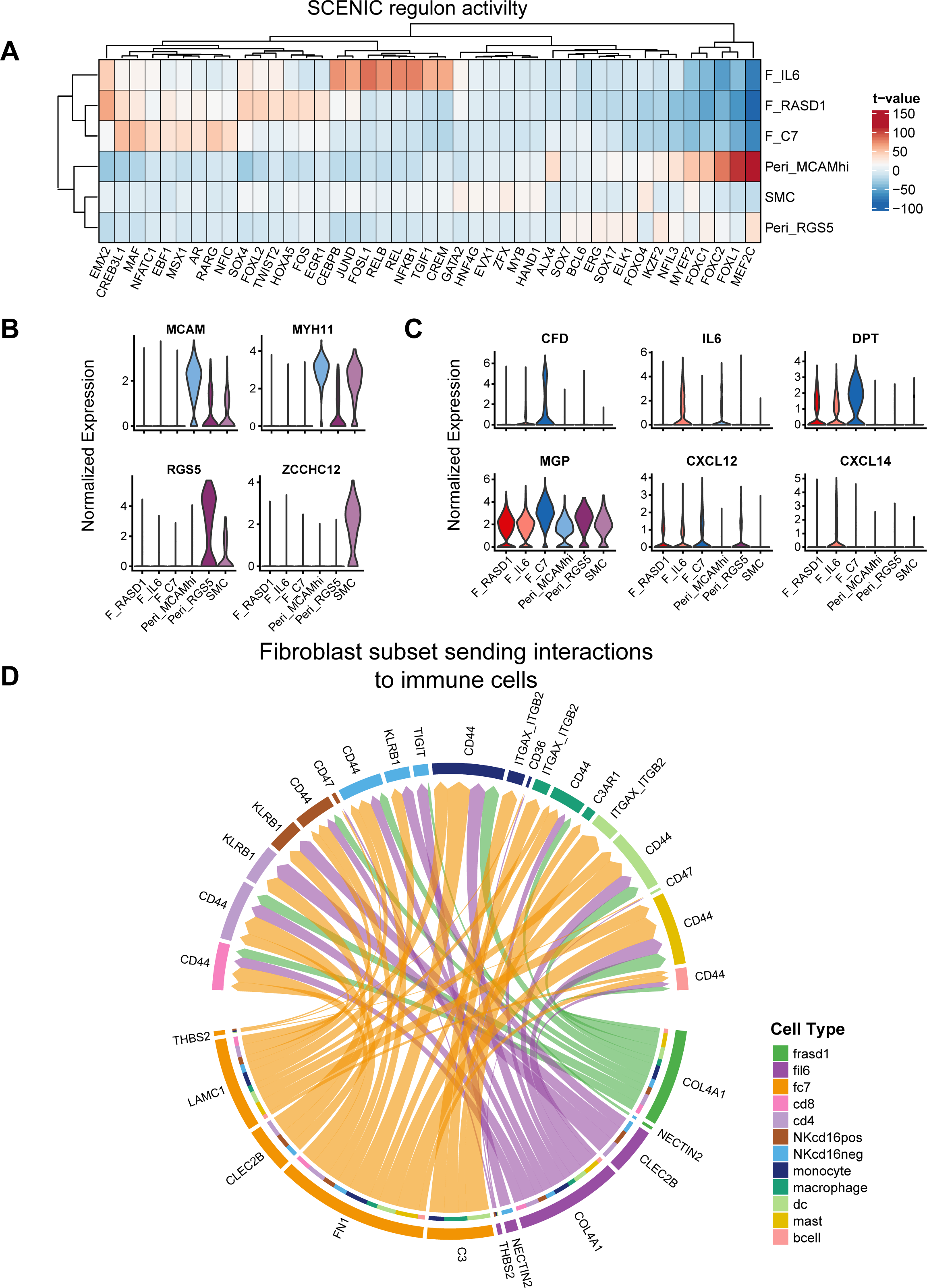
A. Linear modes of SCENIC’s AUC regulon scores by stromal subset, values plotted are extracted t-values from each model. B. Gene signatures of smooth muscle cell and pericyte subsets, values shown are normalized expression values. C. Inflammatory cancer-associated fibroblast gene signatures (Hornburg et al. 2021) in stromal subsets of benign human FTs, normalized gene expression. D. CCIs of fibroblast subsets to immune cells not shared between F_IL6, F_RASD1, and F_C7 clusters. The width of the arrows represents scaled CellChat probability scores.

